# Beyond the Delay Neural Dynamics: a Decoding Strategy for Working Memory Error Reduction

**DOI:** 10.1101/2022.06.01.494426

**Authors:** Zeyuan Ye, Haoran Li, Liang Tian, Changsong Zhou

## Abstract

Understanding how the brain preserves information despite intrinsic noise is a fundamental question in working memory. Typical working memory tasks consist of delay phase for maintaining information, and decoding phase for retrieving information. While previous works have focused on the delay neural dynamics, it is poorly understood whether and how the neural process during decoding phase reduces memory error. We studied this question by training recurrent neural networks (RNNs) on a color delayed-response task. We found that the trained RNNs reduce the memory error of high-probability-occurring colors (common colors) by decoding/attributing a broader range of neural states to them during decoding phase. This decoding strategy can be further explained by a continuing converging neural dynamics following delay phase and a non-dynamic biased readout process. Our findings highlight the role of the decoding phase in working memory, suggesting that neural systems deploy multiple strategies across different phases to reduce memory errors.

**Significance:** Preserving information under noise is crucial in working memory. A typical working memory task consists of a delay phase for maintaining information, and a decoding phase for decoding the maintained into an output action. While the delay neural dynamics have been intensively studied, the impact of the decoding phase on memory error reduction remains unexplored. We trained recurrent neural networks (RNNs) on a color delayed-response task and found that RNNs reduce memory error of a color by decoding a larger portion of the neural state to that color. This strategy is supported both by a converging neural dynamic, and a non-dynamic readout process. Our results suggest that neural networks can utilize diverse strategies, beyond delay neural dynamics, to reduce memory errors.

## Introduction

Working memory is the ability to maintain information for a short period of time without external stimuli. It largely consists of a perception phase for sensing the information, a delay phase for maintaining information, and finally, a decoding phase (for example, response epoch in typical delayed-response tasks^1–3)^ for retrieving information^4^. Working memory tasks are fundamentally challenging because the neural system needs to maintain accurate information despite intrinsic stochastic noise. Without additional error-correcting mechanisms, the neural population activity will deviate from its original state, leading to large memory errors^5–8^. The neural mechanisms utilized by the neural system to mitigate these memory errors remain unclear.

Previous works have primarily focused on the neural dynamics during the delay phase^1,5–14^. For instance, the neural system can form discretized attractors to represent some information values^1,9–11^. An attractor is a special neural population state; any small deviation from the attractor state will be brought back. This unique property of attractors can stabilize neural population states against random deviations due to noise, thereby reducing memory errors. This attractor-based memory error correcting mechanism has been supported by experimental behavior data^1^, experimental neural recordings^15^, and simulations from artificial neural networks^7,11,16^. It can also be implemented by artificial neural networks using biologically plausible synaptic rules^9,17^.

However, despite also being a key phase, the role of the decoding phase has largely been neglected. From the information processing perspective, the decoding phase acts as a decoding mapping, which maps the maintained delayed neural state to an output action^18–23^. A change in the decoding mapping will lead to a significant change of behavioral performance. This importance of decoding mapping can be demonstrated in the Brain-Computer Interface (BCI) experiments^23,24^, where the BCI functions as a decoder, mapping neural population states to external actions (e.g. cursor movement). Alterations in the BCI’s decoding mapping would lead to significant action errors, necessitating relearning of the new decoding pattern by the animal^23,24^. Similarly, in working memory, the delay neural population state must progress through decoding phase, be decoded to muscles signals for an output action (e.g. saccade). What is the decoding mapping from an end-of--delay neural state to an output information? Whether and how such mapping helps reducing the memory error? What processes occur during the decoding phase to establish this decoding mapping? Addressing these questions is important for understanding the diverse strategies, beyond just delay neural dynamics, of the neural system to reduce the memory error.

In this paper, we trained artificial recurrent neural networks (RNNs) to perform a color delayed-response task, where the color in each trial was sampled from a prior distribution with a few high-probability colors (common colors)^11,25,26^. We found that the trained RNN exhibited smaller memory error on common colors, which aligns with previous behavioral experiments^1^. We found two main mechanisms that the RNNs used to reduce memory error. First, the neural system created attractors to encode common colors during the delay phase. Second, during the decoding phase, a large part of the neural population states was decoded to common colors, which improved noise tolerance. This noise-tolerant decoding mapping can be further understood by (1) continuing the attractor-based dynamics during the decoding phase, and (2) a non-dynamic biased readout from the recurrent neural state to common colors. Further, we proposed an approximation formula which naturally decompose the memory error into delay dynamic and decoding components – neglecting decoding component will lead to a failure of explaining the RNN’s memory error. Our results emphasize the importance of the decoding phase and propose experimentally testable predictions.

## Results

### RNNs trained in a biased environment reproduce human behavioural features in the color delayed-response task

Our study utilized a vanilla RNN architecture, which included 12 perception neurons, one go neuron, 256 recurrent neurons, and 12 response neurons (Figure 1A and Methods). We added noise to the recurrent neurons to simulate intrinsic brain noise. Task consists of a fixation, a perception, a delay, a go cue and finally a response epoch^1^. Specifically, in each trial, we sampled a color from an environmental prior distribution with four peaks corresponding to common colors (similar to the behavioral experiment^1^). The sampled input color was perceived by the perception neurons through predefined tuning curves (von Mises functions with shifting mean positions, see Figure 1A), and we added Gaussian noise perturbations to simulate perception noise (see Methods). The perception neurons then fed the activity forward to the 256 recurrent neurons. During the delay period (randomly determined delay epoch duration), the RNN used these recurrent neurons to maintain the color information. After the delay, the go neuron received a signal that initiated the response action. The response neurons, equal in number to the perception neurons, activated during the response epoch, were expected to mirror the activity of the perception neurons during the perception epoch (see Methods). We then averaged the activity of these response neurons over time and mapped it back into color information using the population vector method. Go and response epochs in combination is considered as a decoding phase in this paper.

**Fig 1.**
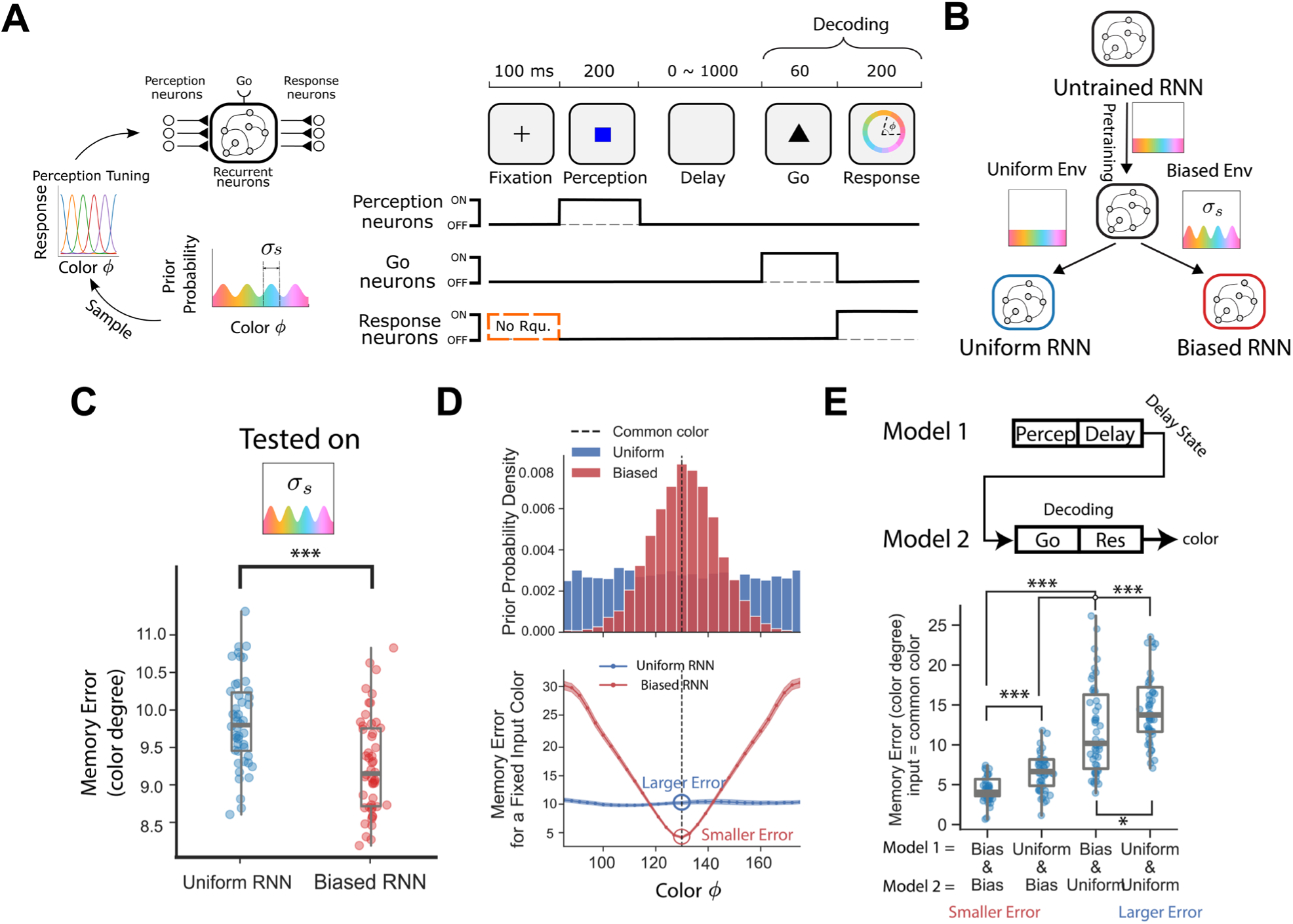
Decoding phase (go and response epochs) is crucial for RNNs to reduce memory error. (**A**) An environment was defined as a probability prior function for the input color. In each trial, a color was sampled from the prior (four common colors = 40^∘^, 130^∘^, 220^∘^, 310^∘^), then sensed by the perception neurons. Following the perception epoch, the RNN went through the delay epoch for maintain information (delay length was sampled randomly from 0 to 1000 ms), go epoch to be ready, and finally response epoch to reproduce the previously sensed color. Except regularization terms, loss function was only applied on the response neurons, which had no requirement in fixation epoch (No Rqu.) and should be silent in the perception, delay and go epochs. (**B**) An RNN was trained by multiple trials progressively (see texts and Methods). Depending on the retrained environment prior, the final trained RNN was categorized as a Uniform RNN (uniform prior) or a Biased RNN (biased prior). **(C)** After training, RNNs were evaluated under a biased environment (σ_*s*_= 12.5^∘^, same as the training environment of the Biased RNN). The memory error of each RNN was calculated as the root-mean-squared difference between output and input colors over 5000 trials. Each dot represents one RNN, with a total of 50 RNNs used for each type. (***: *p* < 10^−3^ Wilcoxon signed-rank test). Outlier RNNs were not shown. **(D)** Top: The prior distribution of colors around a common color (130^∘^, black dashed line). Bottom: Memory error for various input colors, assessed over 5000 trials for each input color and RNN. The colored lines represent the mean memory errors across the 50 RNNs, with error bands indicating standard errors. Outlier RNNs were removed (see Methods). **(E)** Cross decoding suggests the importance of decoding phase in reducing memory error (e.g., Bias & Bias vs. Bias & Uniform). Delay recurrent neural state was prepared from a model 1, then was decoded by model 2. Each dot is one randomly sampled model1-model2 pair. 50 pairs were sampled for each model1-model2 category combination. ***: *p* < 10^−3^; *: *p* < 5 × 10^−2^; Wilcoxon signed-rank test. Boxes indicate the interquartile range between the first and third quartiles with the central mark inside each box indicating the median. Whiskers extend to the lowest and highest values within 1.5 times the interquartile range. Outlier RNNs were not shown. Biased RNN was trained under σ_*s*_ = 12.5^∘^ in panel (C, D, E).

Each RNN was trained progressively (Figure 1B and Method)^27^. The progressive training consisted of two stages. First, in the pretraining stage, the randomly initialized RNN was trained in a uniform-prior environment. Following this, the pretrained RNN was retrained in a new environment. If the new environmental prior is biased, the retrained RNN is referred to as the “Biased RNN”. Conversely, if the new environmental prior is uniform, the trained RNN is called the “Uniform RNN”. Our procedure ensured that both types of RNNs had the same number of training trials.

Similar training with biased environments has been performed experimentally before (with humans)^1,28^. We then asked whether our Biased RNN could reproduce some basic behavioral features of those experiments. We found that (see SI Figure 1), in alignment with previous experiments^1^, (1) The RNN’s memory error increases with longer delays; (2) Despite using uniformly sampled colors as inputs, Biased RNN still outputs biased color distribution, with four output color distribution peaks aligned with the four biased environmental priors in the training; (3) When the input trial’s color is near the common color, the output color of the Biased RNN tends to shift closer to the common colors, suggesting that Biased RNNs have a tendency to output common colors.

### Cross-decoding experiments showed that both delay and decoding phases are crucial in memory error reduction

Using the trained Biased and Uniform RNNs, we then explored the properties of memory error. Memory error of a trial is defined as the circular subtraction (circular period is 360^∘^) of the RNN’s output color from the trial’s input color. The memory error across multiple trials is defined as the root-mean-square error of individual trial errors.

First, we evaluated the RNN’s overall memory error in a biased environment (σ_*s*_ = 12.5^∘^) which is the same environment the Biased RNNs were trained in. As expected, the Biased RNN exhibited smaller memory errors (Figure 1C, *p* < 10^−3^, Wilcoxon signed-rank test). Mathematically in the limit of large number of trials, the memory error is the weighted average (weighted by the prior) of the memory error for every fixed input color. Hence, we examined the memory error for each fixed input color (Figure 1D). For the Uniform RNN, memory errors were uniform across all input colors. In contrast, the Biased RNN showed a reduced memory error for the common color (e.g., at 130^∘^) compared to other input colors and also to the same input color in the Uniform RNN. This reduction of memory error on common colors provided an opportunity to explore the RNN’s neural mechanisms underlying memory error reduction.

Previous studies have primarily focused on delay dynamic mechanisms to reduce memory errors^1^. We hypothesized that, in addition to neural dynamics during the delay, the decoding from the end-of-delay neural state to an output color during decoding phase is also crucial. To test this, we conducted a cross-decoding experiment, where the non-decoding (fixation, perception, and delay epochs) and decoding phases (go and response epochs) were separated (see Figure 1E, upper panel). In this experiment, two RNN models were randomly chosen from either the Biased or Uniform category (two models can belong to the same or different categories). Model 1 underwent a trial with a fixed common color input, and the recurrent neural state (i.e. recurrent neural population activity, or neural state for short) at the delay’s end was collected. Collected recurrent neural state was then used to set the recurrent neural state of model 2. This step requires a one-to-one matching from model 1’s recurrent neurons to model 2’s recurrent neurons. This matching was achieved by comparing the preferred colors of the individual neurons between the two neural populations (see Methods). Model 2 then directly ran through the decoding phase (go and response epochs) to output a color. The error between the output and input color across 500 trials was the memory error of this single RNN pairing (model 1-model 2). This entire process was conducted for 50 random pairs for each category combination (Biased & Uniform, Biased & Biased, Uniform & Uniform, Uniform & Biased). It should be noted that the algorithm used for matching (comparing preferred colors) from the recurrent neural state of model 1 to model 2 is not optimal because the connections are not exactly the same between RNN models. However, we applied the same matching algorithm to all category combinations, so comparisons between different category combinations should be fair to a certain extent.

We found that (Figure 1E), for the same type of delay state preparation RNNs (model 1), decoding using Biased RNNs significantly reduced memory error of common colors compared to decoding using Uniform RNNs (i.e., Bias & Bias vs. Bias & Uniform, and Uniform & Bias vs. Uniform & Uniform). This suggests that Biased RNNs learned certain neural mechanisms during the decoding phase to reduce the memory error of common colors.

Besides, we found that for a fixed type of decoding RNN (model 2), using Biased RNN for delay state preparation also results in smaller memory errors of common colors. This finding supports the idea that Biased RNN learned neural mechanisms during the delay phase to reduce memory errors.

### A hypothesis of the memory error reduction by both delay neural dynamics and decoding strategy

The cross-decoding experiments conducted underscore the pivotal roles of both the delay and decoding phases (go cue and response epochs) in influencing memory errors. Previous hypotheses have suggested that reduced neural dispersion dynamics during the delay might contribute to error diminution. In this paper, we introduced a novel hypothesis concerning the importance of the decoding phase in error reduction. Conceptually, the state of a neural population can be represented as a point within a multidimensional state space, where each axis represents the activity of an individual neuron (Figure 2). The end-to-end effect of decoding phase is to transform the neural state at the end of the delay into an output color. Geometrically, decoding phase “maps” distinct potions in the state space to specific output colors. Imagine a scenario where neural states, despite being noisy and dispersed, fall within a region of the state space mapped to a single color; the output color remains constant despite the dispersion of neural states. Thus, colors with larger state space occupancies exhibit greater tolerance to noise, subsequently diminishing memory errors. In summary, we propose that memory error reduction results from two factors (Figure 2): (1) reducing the dynamic dispersion of neural population states during the delay, and (2) assigning a broader range of neural states to common colors during the decoding phase, a process we term ‘larger state space occupancy by a color’. This paper primarily tests and explores the second mechanism, focusing on the effect of decoding phase on memory error reduction.

**Figure 2.**
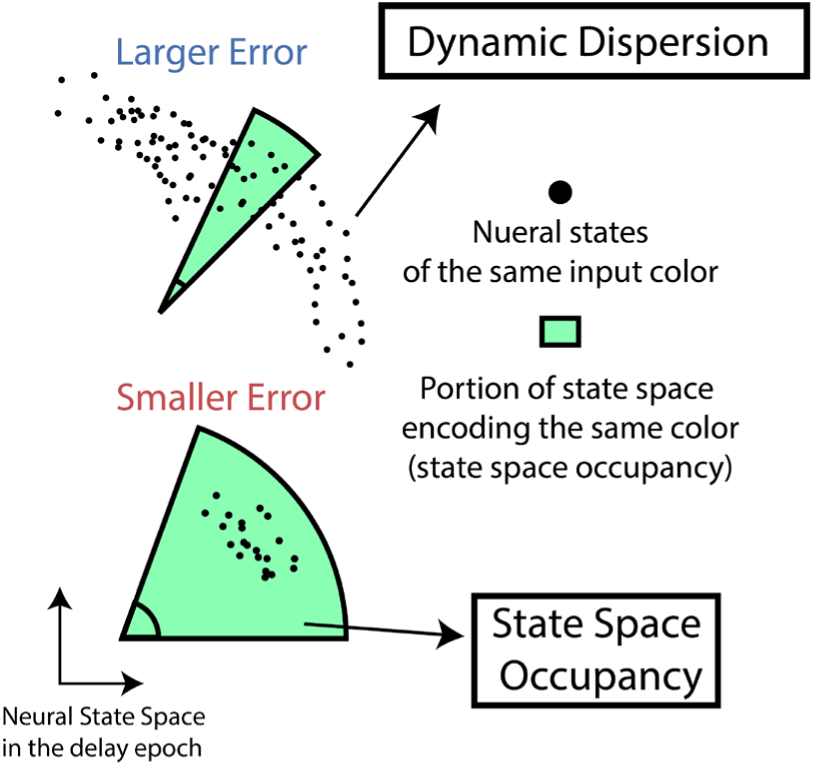
A hypothesis decomposing the memory error reduction mechanism into delay neural dynamics (dynamic dispersion) and a decoding strategy (state space occupancy). Illustration of the hypothesis: (1) Each axis is one recurrent neural activity. Each dot is one recurrent neural population state at the end of the delay under a fixed input color. Memory error can be reduced if the neural states are more stable (smaller dynamic dispersion). (2) Green region represents portion of state space that will be decoded into a color. Larger state space occupancy of one color allows better tolerance of neural state dispersion, hence smaller memory error for that color.

### RNN’s neural activity is low-dimensional

To test our hypothesis on memory error reduction, we inspected the neural activities in the RNNs. We conducted multiple trials for each RNN (intrinsic noise was turned off), using uniformly sampled color inputs. Recurrent neural activities were collected and projected onto the first few principal components (PCs) using Principal Component Analysis (PCA). The cumulative variance explained ratio suggests that the neural population activity is essentially low-dimensional (Figure 3A): requiring 3 dimensions to account for over 90% of the variance in neural activity across the entire trial, and only 2 dimensions when focusing exclusively on the delay epoch.

**Figure 3.**
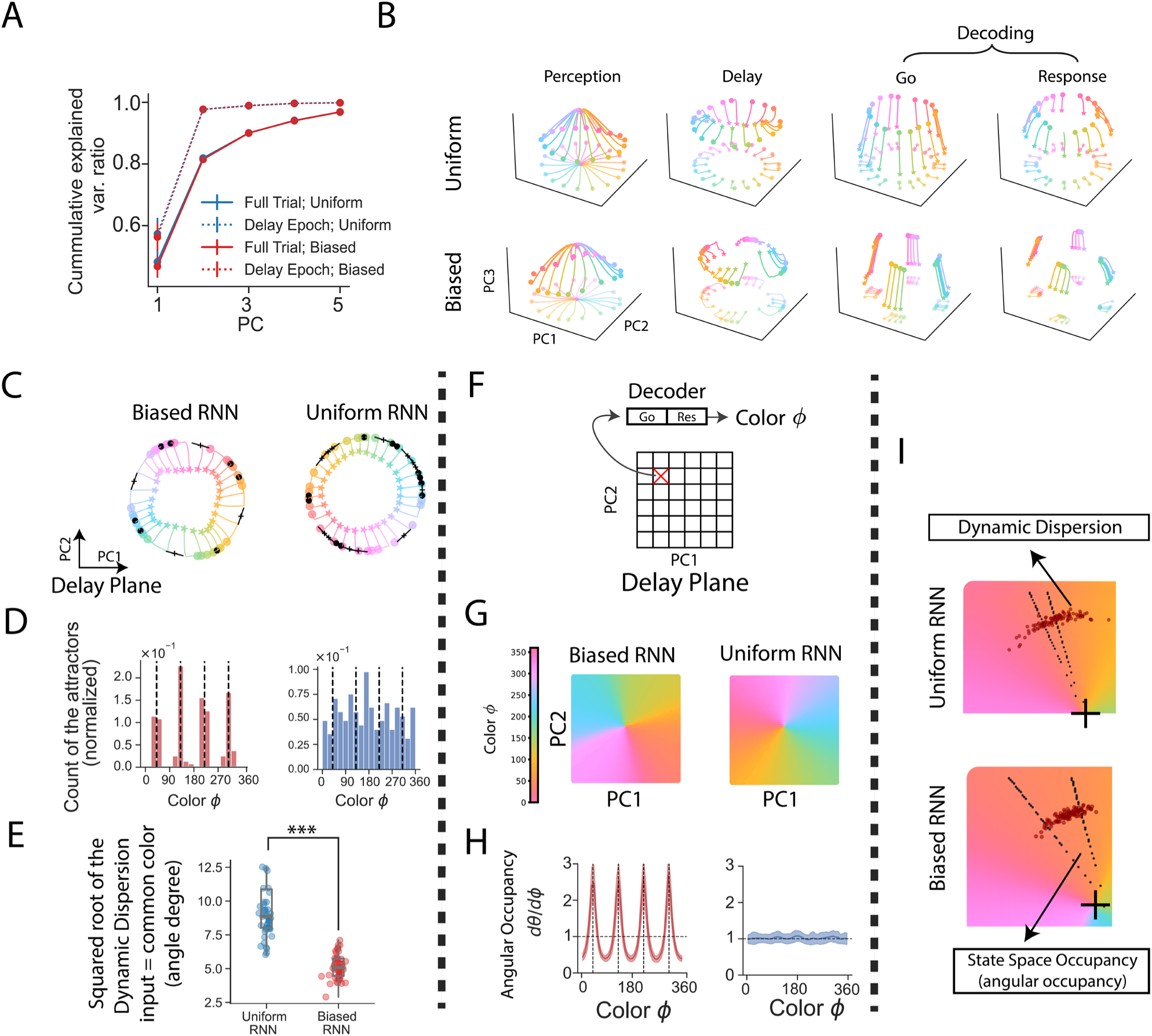
Reverse engineering of the RNNs shows evidence for the hypothesized memory reduction mechanisms (smaller dynamic dispersion and larger state space occupancy). Biased RNN was trained under *σ*_*s*_ = 12.5^∘^. **(A)** Dimensionality of the neural activity (1000 trials for each RNN). Dots and error bar represent mean and standard deviation across 50 RNNs. Uniform: Uniform RNN; Biased: Biased RNN. Blue and red traces mostly overlap. **(B)** PCA visualization of neural states. Stars denote the beginning of the epochs; dots represent the end. Each trajectory is one trial with color indicating the trial input color. Both 3D trajectories (upper) and their 2D projections (lower) were shown in each panel. **(C)** 2D visualization of two example RNN’s neural states in the delay epoch. Black dots are the attractor points. Crosses are saddle points whose long bars indicating the unstable (positive eigenvalue) directions. **(D)** Distribution of attractors. More attractors were formed near common colors (dash lines) in the trained Biased RNN. Results concatenated attractors from 50 RNNs. **(E)** End-of-delay neural states have smaller dynamic dispersion (see Methods) in Biased RNN when the input color is fixed to common color. Each dot is one RNN. ***: *p* < 0.001 Wilcoxon signed-rank test. Boxes indicate the interquartile range between the first and third quartiles with the central mark inside each box indicating the median. Whiskers extend to the lowest and highest values within 1.5 times the interquartile range. Outlier RNNs were not shown. **(F)** Decoding the delay plane by selecting mesh points and running through go and response epochs. **(G)** Decoding results of two example RNNs. Color is the decoded color. **(H)** Angular occupancy--the size of angle (θ) on the delay plane used for encoding one unit of color. Solid lines are the mean across 50 RNNs, and error band is the standard deviation. Outlier RNNs were removed (see Methods). (**I**) Two example RNNs illustrate the joint effect of dynamic dispersion and angular occupancy. Each black dot is the position of neural state (on the delay PC1-PC2 plane) at the end of delay of one trial (delay length = 800 ms), where trial’s input color was fixed to a common color (40^∘^). Dashed lines enclose the angular space for encoding 35^∘^ to 45^∘^. Black crosses are the center of the PC1-PC2 plane. In this figure, RNN’s intrinsic noise was turned off for better visualization, except in (E, I).

The inherent low dimensionality of the neural activity allowed for direct visualization by projecting the activity onto the first three PCs, as shown in Figure 3B. During the perception epoch, the neural states are driven by the input colors, moving towards different directions within a ring-shaped manifold. In the subsequent delay phase, these neural states remain relatively stable, although slight lateral movements are observed. Next, in the go epoch, the neural state shifts along the third principal component. Finally, in the response epoch, the neural states roughly stay stable in a 2D ring-like structure. These observations from the various phases of neural activity illustrate the dynamic yet structured nature of neural state dynamics in different phases of task execution.

### Biased RNN formed attractors near common colors to reduce the dynamic dispersion

Next, we tested whether the dynamic dispersion during the delay phase is a key factor in memory error reduction. We projected the delay neural activity to the first two PCs. We refer this 2D representation as the “delay plane”. On the delay plane, neural states for different input colors formed a ring-like structure (Figure 3C).

We observed small lateral motion of neural states along the ring. According to the dynamic theory, this small motion may be driven by fixed points on the ring. To test this, we searched fixed points by finding the local minimum of speed (see Methods). We found two types of fixed points on the ring: (1) Attractors which attract nearby states (2) Saddle points, which repel nearby neural states along the direction of positive eigenvalues. Interestingly, we found in the Biased RNN, attractors predominantly appeared in four positions (Figure 3C). These attractors also had more negative eigenvalues (indicating stronger attractive force) in more Biased RNNs (with smaller prior σ_*s*_, SI Figure 2).

The four dominant positions of four attractors may represent the four common colors in the biased environmental prior. To test this, we developed an RNN decoder. The RNN decoder itself is a faithful copy of the original RNN. Given a neural state of interest, the RNN decoder’s recurrent state is set to match the neural state, then without any delay, it directly runs through the go and response epoch. The output color is the decoded color of the neural state of interest.

We decoded the attractors of Uniform RNNs and Biased RNNs. The results revealed that in Biased RNNs, attractors were primarily mapped to common colors (Figure 3D). Since attractors have “attracting effects,” this suggests that the neural state may have a smaller dynamic dispersion when the input color is a common color. To test this, we directly measured dynamic dispersion as follows: We conducted multiple trials for each RNN, keeping the input color fixed at the common color and the fixed the delay length at 800 ms. Neural states at the end of the delay were collected, and their angles on the delay plane were computed. The variance of these neural state angles represents the dynamic dispersion. The results (Figure 3E) indicate that the dynamic dispersion for the common color in Biased RNNs is smaller than in Uniform RNNs, supporting the hypothesis that RNNs reduce memory error by diminishing dynamic dispersion^10^.

### Biased RNN allocates larger angular occupancies to common colors

Next, we tested whether the RNNs formed larger state space occupancy to common colors. The end-to-end effect of decoding phase is transforming neural state at the end of delay epoch to an output color. This end-to-end effect is entirely same as the RNN decoder (Figure 3F), which decodes a neural state by continuing to run the neural state through go and response epochs. We used the RNN decoder to decode the delay plane (not the neural state from any particular trials). Example RNNs’ results suggested that color information is represented as angles in the delay plane (Figure 3G).

We investigated the quantitative relationship between angle and color. We sampled and decoded dense neural states along a ring (see Methods). This provided a numerical relation between the angle to the decoded color. The numerical differentiation of the angle by the color is called angular occupancy. Angular occupancy measures the size of angular space mapped to a unit change of color. Results indicate that, in the Biased RNN, common color has larger angular occupancy (Figure 3H). As a baseline comparison, the angular occupancy of the Uniform RNN is uniform. These results suggested that Biased RNN reduces the memory error (for common color) by allocating larger angular space.

We used two example RNNs to provide an intuition of how the dynamics and decoding strategy jointly contribute to the reduction of memory error (Figure 3I). We ran each of the example RNNs through 1000 trials with a fixed common color as input (delay length fixed at 800 ms). Neural states at the end of the delay were collected and visualized on the delay plane. Two dashed lines enclosed the portion of space encoding color 35^∘^to 45^∘^, around a common color 40^∘^. It can be observed that, compared to the Uniform RNN, the Biased RNN has a larger angular occupancy and smaller dynamic dispersion, supporting the hypothetical picture in Figure 2.

### An approximate theory quantitatively relates the dynamic dispersion and angular occupancy to the memory error

Figure 3I provides a visualization of how the dynamic and decoding strategy jointly contribute to the reduction of memory errors on common colors. Here we study a mathematical conceptualization of how dynamic dispersion and angular occupancy jointly contribute to the reduction of memory errors.

The central idea is the Taylor expansion (see Methods). Color *ϕ* is a function of angle *θ* in PC1-PC2 space. Denoting common color as *ϕ*_*c*_, the corresponding angle as *θ*_*c*_. Assuming that, when trial input is the common color, the actual angle of neural state *θ* does not deviate from the true color angle *θ*_*c*_ too much. Therefore, we expanded the color function as *ϕ*(*θ*) ≈ *ϕ*_*c*_ + (*dϕ*/*dθ*|_*ϕc*_)(*θ* − *θ*_*c*_), where *dϕ*/*dθ*|_*ϕc*_ is the reciprocal of angular occupancy. Inserting this approximation into the squared memory error 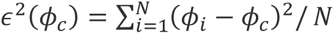 (where *ϕ*_*i*_ is the output color in trial *i*), we can then obtain an approximation formula (Equation 1 and Figure 4A) describing the memory error using angular occupancy (*dϕ*/*dθ*|_*ϕc*_), dynamic dispersion 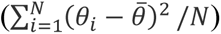 and a mean bias correction ((*θ̅* − *θ*_*c*_)^2^)

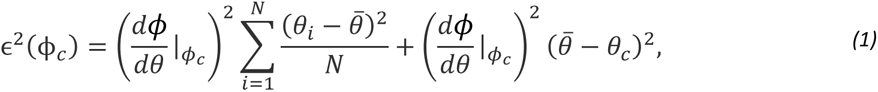

where the mean bias correction described the correction due to the shift of trial averaged mean angle *θ̅* to *θ*_*c*_.

**Figure 4.**
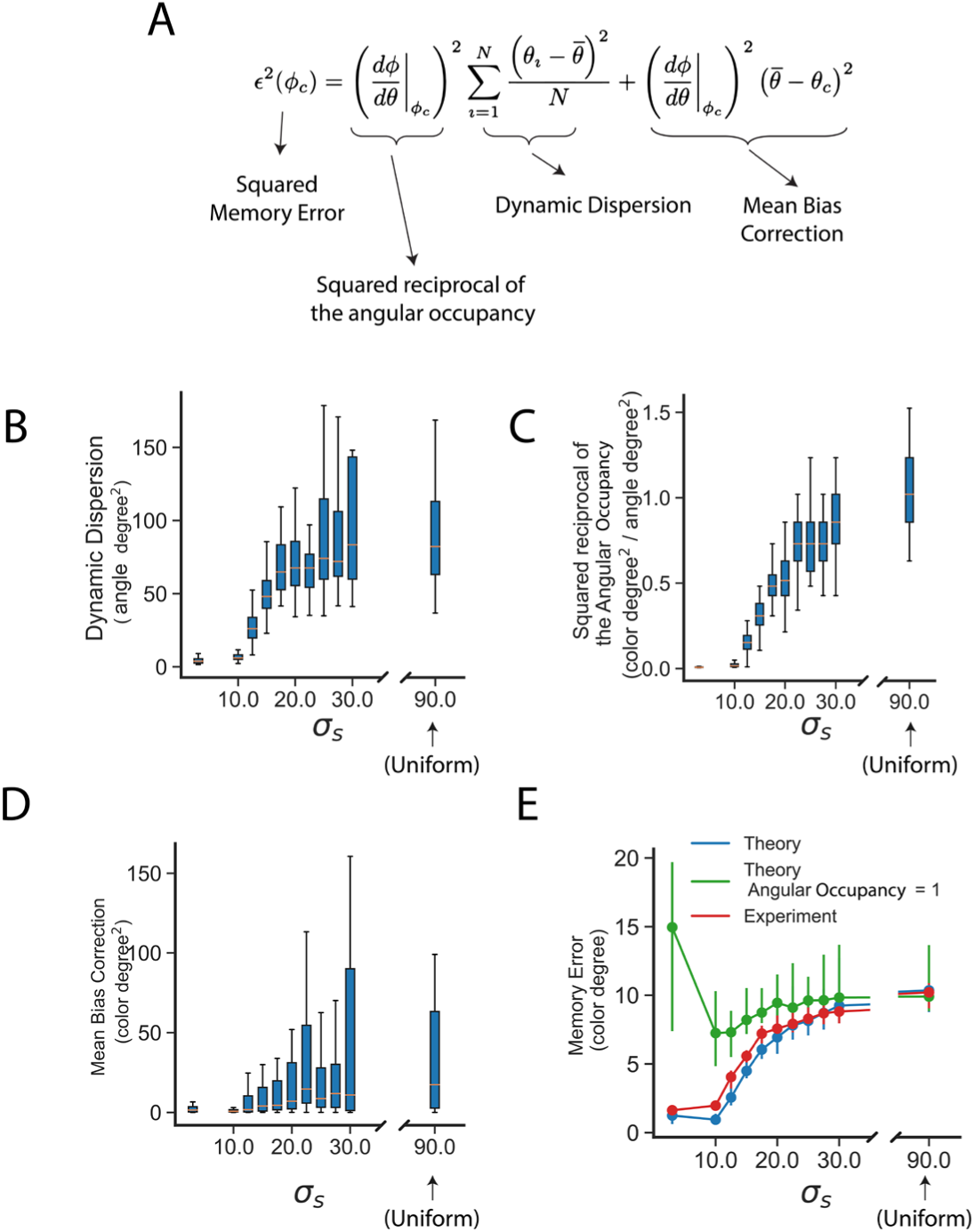
An approximation formula decomposes the memory error of a common color into a dynamic dispersion, angular occupancy, and a correction term. **(A)** An approximation formula for the squared memory error (see text). **(B, C, D)** Dynamic dispersions, squared reciprocal of the angular occupancy and mean bias correction for biased RNNs trained on different biased environment (50 RNNs for each σ_*s*_). Smaller σ_*s*_ indicating narrower prior (see Figure 1A). Boxes indicate the interquartile range between the first and third quartiles cross 50 RNNs with the central mark inside each box indicating the median. Whiskers extend to the lowest and highest values within 1.5 times the interquartile range. **(E)** Comparing the theory to the experimental RNN’s memory error. Dots are the median memory error, and error bars show first and the third quantiles across 50 RNNs for each σ_*s*_. Theoretical prediction was computed as (A). Experimental memory error of each RNN was measured by computing the averaged memory error of 5000 trials (fixing input at common color, outlier trial errors were removed). As a comparison, we also computed the prediction of theory but assuming a trivial angular occupancy (equals to 1).

To test this approximation formula (Figure 4A), we trained biased RNNs on different environmental priors by changing the value of *σ*_*s*_ to control the prior width. The smaller the *σ*_*s*_, the narrower the prior, and the higher the prior probability of sampling a common color. We then computed dynamic dispersion, squared reciprocal of the angular occupancy and mean biased correction separately (see Methods). On the other hand, we also computed the experimental memory error directly by running each RNN 5000 trials, fixing input color as common color.

We found the higher common color prior probability is (smaller σ_*s*_), the smaller dynamic dispersion and the larger angular occupancy (Figure 4B, C) are. In general, the theoretical prediction has good alignment with the actual experimental memory error (Figure 4E). To test whether the angular occupancy term is important for the memory error, we computed theoretical prediction (Figure 4A) but setting angular occupancy to 1. We showed that neglecting angular occupancy factor will lead to a failure of explaining the RNN’s memory error for small σ_*s*_ (Figure 4E).

### Higher intrinsic noise leads to larger angular occupancy for common colors

This approximate theory is general in principle. It not only works for different environment priors, but in principle should also work for the same environmental prior with different RNN’s noise strength (Figure 5). This scenario potentially mimics brains with different noise level.

**Figure 5.**
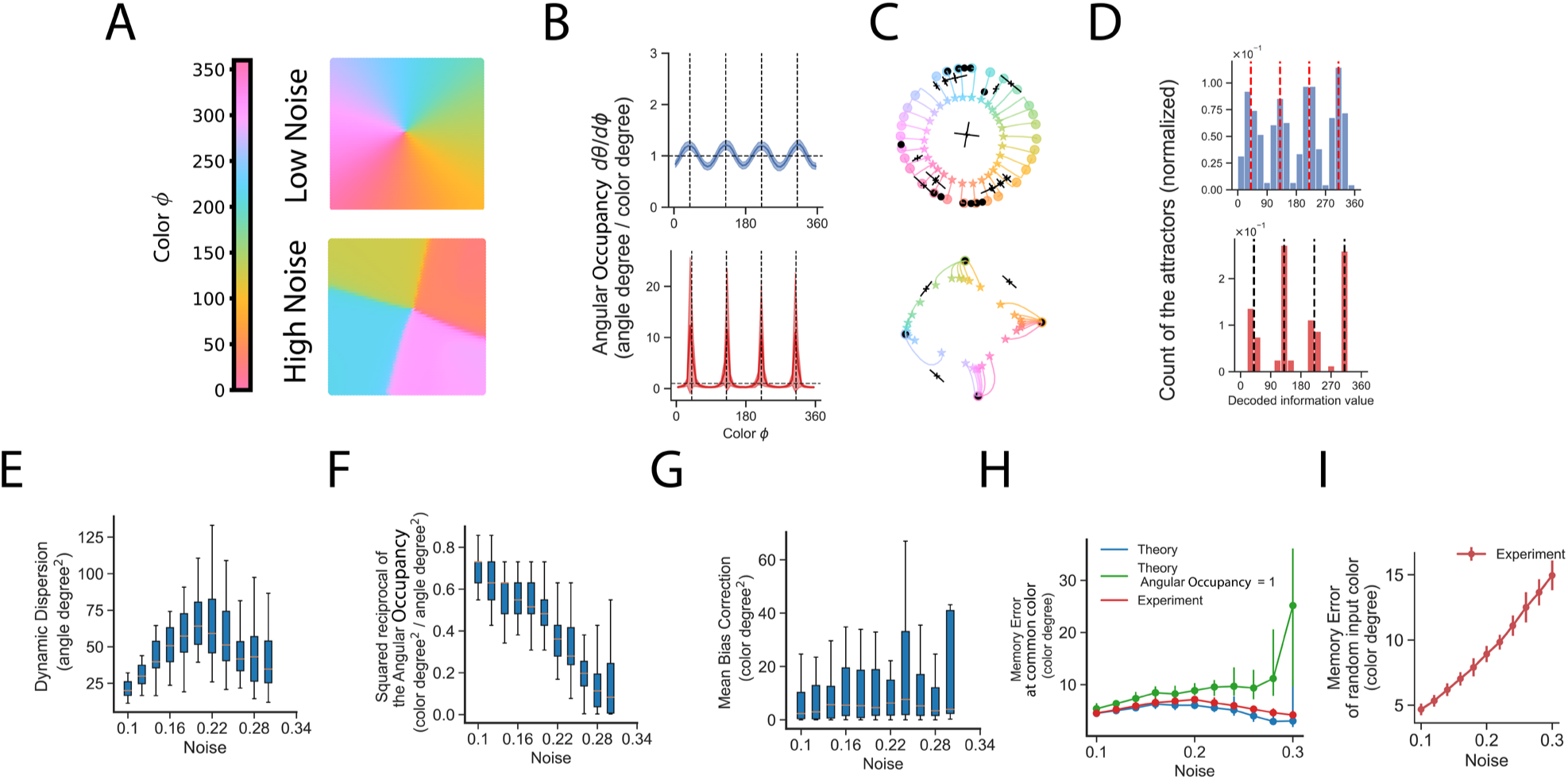
The results for different noise strength also support that angular occupancy is an important factor to reduce memory error at common color. (**A**) Decoding the delay plane for two example RNNs trained on low-noise (noise strength σ*_rec_* = 0.1) and high noise (σ*_rec_* = 0.3). (**B**) Angular occupancy. Dash vertical lines are common colors. Solid blue/red line indicates the mean of 50 RNNs and error bands are standard deviation. (**C**) Neural dynamics during delay. Each trajectory is the delay of one trial with trajectory color indicating the input color. Stars are the beginning of delay; dots are the ends. Black dots are attractors, black crosses are saddles with long bar indicating the positive eigenvalue directions. (**D**) Number of attractors for different colors. Attractors were concatenated from 50 RNNs. (**E, F, G**) Dynamic dispersion, squared reciprocal of the angular occupancy and mean bias correction measured for 50 RNNs for each of the noise strength (see Methods). Boxes indicate the interquartile range between the first and third quartiles cross 50 RNNs with the central mark inside each box indicating the median. Whiskers extend to the lowest and highest values within 1.5 times the interquartile range. (**H**) Comparing experimental results with theoretical predictions for memory error. Dots are the median across 50 RNNs and error bar shows first and third quantiles. In the experimental part, memory error of each RNN is the average of 5000 trials error fixing input as common color (outlier trials’ error were removed, see Methods). (**I**) Experimental error same as (H), but input colors were randomly sampled from an environmental prior σ_*s*_ = 17.5^∘^ instead of fixing to a common color. In this figure, all RNNs were trained on environmental prior *σ*_*s*_ = 17.5^∘^. Noise was turned off after training in computing panel (A, B, C, D, F).

We trained 50 RNNs under high noise and low noise conditions, respectively. Same procedures were used to compute the RNNs attractors and angular occupancy. We found that high-noise RNNs have larger angular occupancy to common colors (Figure 5A, B), and also more likely to form attractors to represent common colors^1,11^ (Figure 5C, D).

To test whether the approximate formula (Equation 1) still works in this varying noise case, we trained RNNs using different noise levels. Their dynamic dispersion, space occupancy and mean bias correction were computed separately (Figure 5E, F, and G). Experimental memory error was also computed by directly comparing input color to output color. Again, the approximate formula aligns with experimental memory error well, and better than theory without considering the decoding strategy (angular occupancy = 1) (Figure 5H). Note the error studied in Figure 5H is only the error of the common colors. It can be observed that total error averaged across different input colors naturally increases under larger noise conditions (Figure 5I).

### Larger angular occupancy is due to both decoding dynamics and biased readout in the decoding phase

We have shown that the decoding strategy (i.e., angular occupancy to a color) is an important mechanism for reducing memory errors. This is an end-to-end effect of decoding phase. Next, we reverse-engineered the decoding phase to explore what happens inside the decoding phase that led to such a non-uniform angular occupancy. We study this problem by dividing and conquering. The decoding phase can be roughly separated into three stages (Figure 3B, 6A). (1) Go Dynamics: a go cue signal pushes the recurrent neural states from the delay plane to the response plane. (2) Response Dynamics: recurrent neural states can dynamically move during the response epoch (epoch length = 200 ms). (3) Readout: during the response epoch, recurrent neural states were readout (Equation 3 in Methods) into response neural activity, then mapped into output colors (Equation 7). The first two stages can be seen as continuing neural dynamics following the delay epoch (although there is a go cue input during the go epoch, altering the dynamic trajectory). The third stage, readout, does not involve dynamics. It simply multiplies the recurrent neural states with a readout matrix, adds a bias current constant, and then transforms it into output color. Biologically speaking, readout matrix may involve the neural connections from motor cortex to muscles (for actions); bias current constant reflects the intrinsic property of each neuron (e.g. describing the diversity of neural firing threshold). We inspect possible mechanisms leading to non-uniform angular occupancy for each of these three stages (Go Dynamics, Response Dynamics, and Readout) separately.

In the Go Dynamics, tangent motion is a possible mechanism leading to larger angular occupancy for common color. We isolated the go epoch and measured its dynamics. At the beginning, we sampled a uniform ring of neural states (Figure 6B) in the delay plane, then let all these states evolve through go epoch (60 ms). At the end of the go epoch, we collected those neural states and fitted PC1-PC2 plane again. This refitted PC1-PC2 plane is called response plane. We found that neural states accumulated to a few positions. The degree of accumulation can be quantified by measuring the entropy of neural state distribution. The smaller the entropy is, the more likely the neural states tend to accumulate to a few positions. Figure 6C suggests that neural states accumulate happens when the environment prior σ_*s*_ is small. This biased motion during the go epoch is a possible reason leading to a non-uniform angular occupancy in the delay plane. For instance, if all neural states move to the same position during the go epoch, they will all be read out to the same color. This means that, regardless of where the delay neural states were, they will all be decoded into the same color. As a result, the angular occupancy is highly biased.

**Figure 6.**
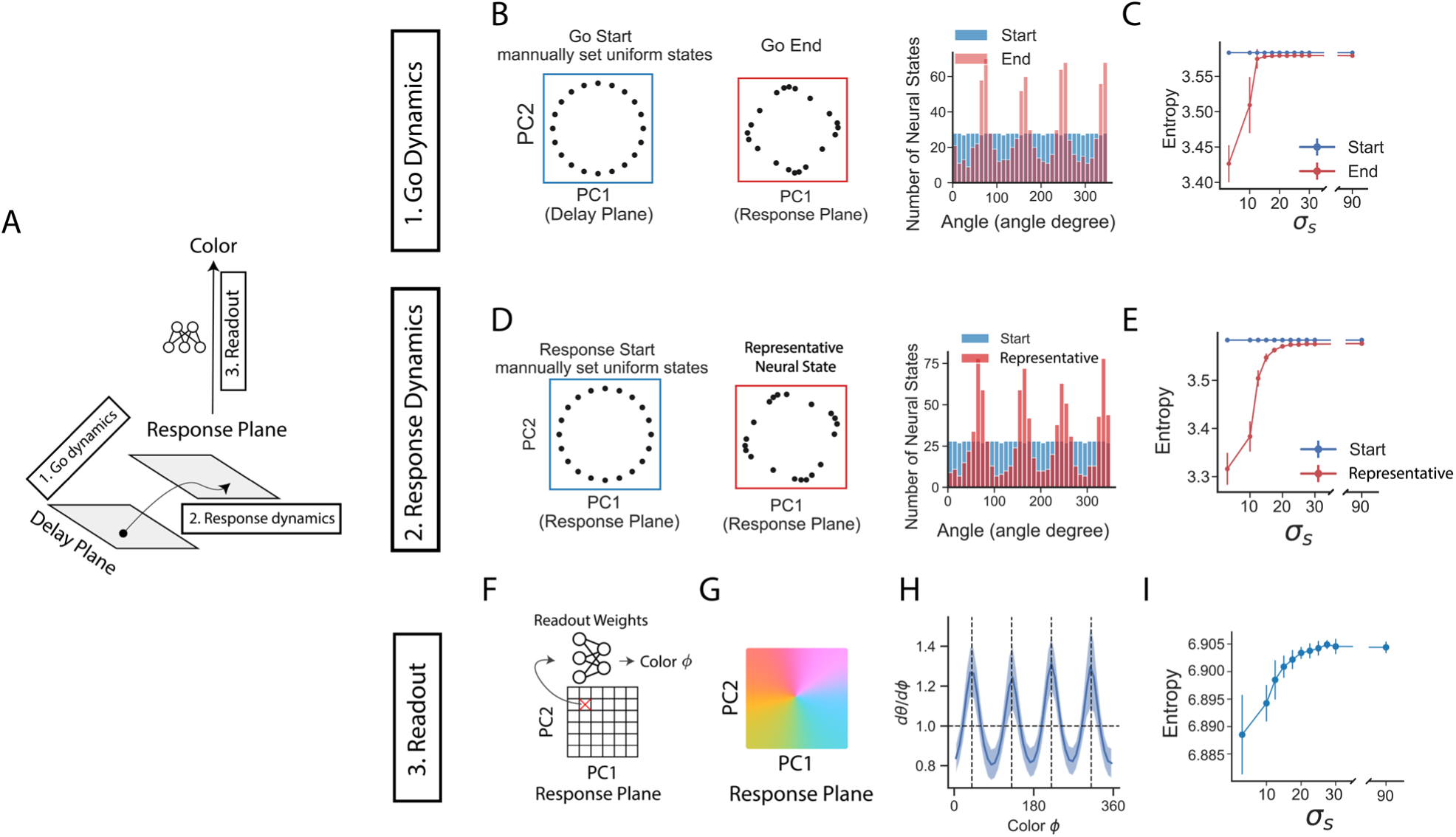
Larger angular occupancy (for common color) is due to both dynamics and biased readout mechanisms during the decoding phase. The single example RNN shown in this figure was trained on prior σ_*s*_ = 3^∘^; delay epoch length is fixed to 800 ms. **(A)** The decoding phase roughly contains three stages (see Figure 3A, B): (1) Go cue drives neural states from delay plane to response plane; (2) Neural dynamics during the response epoch; (3) Readout from recurrent neural states to output color. **(B)** We isolated the effect of Go Dynamics, by firstly fixing neural states uniformly along a ring in the delay plane, then letting the neural states evolve through go epoch. Left two boxes show one example RNN’s neural states at the start and end of Go epoch. Right: the distribution of 1000 neural states of the same example RNN. **(C)** Entropy measures how non-uniform the neural states distribution is. Dots are the mean entropy across 50 RNNs (with outlier RNNs removed), error bar is the standard deviation. **(D, E)** Same as B, C, but the neural states were initialized in the response plane and evolved through response epoch. Representative neural states: recurrent neural states temporally averaged across whole response epoch, which is more relevant for decoding color (see Equation 7). **(F)** Decoding response plane, by selecting mesh points and then readout into color through a readout matrix and bias currents (see Methods, Equation 3 and 7). No dynamics were involved. **(G)** The example RNN’s decoded response planes. **(H)** Angular occupancy in the response plane. Line shows the mean of 50 RNNs (σ_*s*_ = 3^∘^), and error band is the standard deviation. Vertical dash lines: four common colors. Outlier RNNs were removed. **(I)** Entropy of angular occupancy, dots are the mean and error bar are standard deviation (outlier RNNs removed).

Similarly, biased motion during the response epoch may also contribute to non-uniform angular occupancy. As in the study of Go Dynamics, we sampled uniform neural states on a ring in the response plane and allowed them to evolve through the response epoch (200 ms). Since the temporally averaged neural states were used to output a color, we computed the positions of “representative neural states,” which are the temporally averaged neural states during the response epoch (Figure 6D). Results suggested that the tangent motion of neural states also occurs when the environmental prior σ_*s*_ is small (Figure 6E). This implies that angular motion during the response epoch could also be a reason for larger angular occupancy in the delay plane.

On the one hand, the above two dynamic mechanisms are reasonable. The go and response epochs themselves require a duration of time, necessitating the maintenance of information throughout the decoding phase, thus providing an opportunity for neural dynamics to play a role. However, on the other hand, it is important to recognize that the dynamics during the go and response epochs can be quite different from that of delay epoch. The go pulses acts as an input to the neural system, prompting a shift in neural states during the go epoch. This shifted space (response plane) may have different dynamic fields compared to the delay plane^29,30^. Therefore, it should not be regarded as merely a trivial extension of the delay dynamics.

Not only the dynamical evolution of recurrent state during the decoding phase matters, how the recurrent neural state maps to an output color (including the mapping from recurrent neuron to response neuron and population method for mapping response neural activities to an output color, Equation 3 and 7) may be also important. This is called Readout and does not have a time factor hence it is not a dynamics process. As an extreme example, a trivial Readout can map all possible recurrent neural states (in the response plane) into a single color, hence the angular occupancy can be highly biased.

To examine the Readout’s characteristics, we sampled a grid mesh of points within the response plane. Then, utilizing the readout matrix along with the bias current (as elaborated in Equation 3 and the population vector method in Equation 7), we transformed these mesh points into corresponding colors (illustrated in Figure 6G). The resulting data (Figure 6H) revealed that the Readout itself exhibits a bias, mapping larger angular spaces to common colors. This bias becomes increasingly pronounced in a smaller environmental prior σ_*s*_ (Figure 6I).

Overall, these results suggest that angular motion during the go and response epoch, and the biased readout process (including bias current in Equation 3) are the possible reasons for larger angular occupancy for common color in the delay plane. Interestingly, we found that if RNNs were trained on shorter response epochs, response dynamics became weaker while the readout effect became stronger (SI Figure 3). This suggests that dynamic and non-dynamic factors may compensate for each other. It also implies that in some scenarios, where the agent has limited time to respond, the readout mechanism may play a more dominant role in shaping the decoding strategy to reduce memory errors.

## Discussion

Neural states and responses are noisy. How the noisy neural system maintains accurate information is a central question in working memory. Previous researches emphasize neural mechanisms during the delay. Our results provide a novel perspective, highlighting the importance of the decoding phase in reducing memory errors. Smaller memory errors for a color can be achieved by allocating larger state space occupancy (or angular occupancy in this paper). Further analysis revealed that biased state space occupancy is due to both (1) continuing the attractor-based dynamics during the decoding phase (although containing a go cue pulse) and (2) a biased readout from recurrent neural activity to output color. These two factors mutually compensate as shown when training the RNNs with shorter response epoch duration. In general, decoding phase (retrieval of information) is a common phase of working memory tasks, our results encourage certain attention on the role of decoding in diverse working memory tasks.

The logic of this paper resembles a previous human/animal behavioural experiment^1^: training/testing the agent to perform a memory task within a specific prior color distribution, observing a reduction in memory errors for some input colors, and subsequently proposing and/or testing potential neural mechanisms for this error reduction. But we conceive the suggested memory error reduction mechanisms can be more universal than merely adapting to environmental priors. For instance, in Figure 5, we demonstrated that RNNs trained on the same prior but with different noise levels exhibit similar mechanisms for reducing memory errors. In a broader context, mechanisms for reducing memory errors might arise when there is a high demand for accurate retention of information. One potential example is reinforcement learning^31^. The importance of information values is weighted by their associated rewards. Consequently, we further hypothesize that RNNs may adopt similar mechanisms for reducing memory errors (attractor dynamics and a larger state space occupancy) for high reward values. This hypothesis in future can possibly be tested in physiology experiments ^1,30,32,33^ by appropriately assigning rewards to different information values while training animals to perform delayed-response tasks, or can possibly be tested on RNN models using reinforcement learning^34,35^.

Concretely, this paper proposes several experimental testable phenomena. First, when the input value is important (i.e., high-prior color or high-reward value), the neural population dispersion (measured by the variance of repeated trials) should be smaller. Second, state space occupancy for important values should be larger. The state space occupancy can be measured similar to the numerical procedure described here. Specifically, test the animal on multiple trials with different input colors and collect the neural population states at the end of the delay. These neural states, along with the animal’s output color, provide a mapping from neural state space in the delay to the output color. This mapping is determined by the decoding phase hence can be used for studying the decoding strategy. With the mapping from neural state space to output color, numerical differentiation can then be computed. State space occupancy for important colors (high-prior or high-reward) can be compared with other colors. Finally, the approximate formula (Figure 4A-E) can also be tested. For example, train the animal to perform delayed-response tasks in different environmental priors^1,36^, and then measure the dynamic dispersion, angular occupancy, and mean bias correction separately. Theoretical predictions can be compared with the animal’s experimental memory error.

In certain physiology experiments, the neural population state resides within a low-dimensional manifold and employs a ring-like structure to represent information ^13,30,37^, akin to the ring structure discovered in our trained RNNs. In such cases, the state space occupancy can be computed by determining the angle of the neural population state. However, we are aware that in many cases, neural manifolds exist in high dimensions, such as a highly curved trajectory within a high-dimensional space^6,8^. In these instances, state space occupancy does not rely on angular occupancy but rather possibly on “arc-length occupancy”. Arc-length occupancy means the arc length used to represent a unit change of information. Measuring the arc-length occupancy of a highly curved trajectory can be challenging. Yet this challenging can be eased by the concepts and methods advanced in the recent geometric framework^38–42^. Various nonlinear dimensional reduction methods^42^ (e.g., CEBRA^43^, UMAP^44^) can project the manifold into a few latent dimensions for better visualization. Additionally, methods like SPUD^45^ or simple kernel smoothing^36^ enable the direct parameterization of a high-dimensional curve by its arc length, facilitating the measurement of arc length occupancy for each information value. In future, it would be interesting to explore the memory error reduction mechanisms in high-dimensional and more complex manifold scenarios.

We modeled a classical color-delayed response task^9,11,13,16,17,46^ that captures the most essential three stages of memory process: perception, maintenance (delay), and decoding. Beyond this classical experimental design, there are some other memory task variants^1,2,30,36,47,48^. Some tasks have no explicit go cues^1–3^. For example, a color wheel can directly appear on the screen during the response epoch to cue responses^1^. This constant visual feedback to the animal may lead to neural dynamics different from those shown in this paper. Furthermore, multiple items can be shown on the screen for the animal to memorize^49^, for example, simultaneously memorizing multiple colors^30^. How do the neural dynamics of multiple items interact with each other? Does the neural system form a disentanglement representation for different items^30,50,51^, or a shared mechanism for correlated memory items^52^? Is it possible that higher-rewarded items are encoded with larger encoding spaces? Overall, the decoding phase (or so-called memory retrieval) exists ubiquitously in all sorts of complex memory tasks. Our paper emphasizes the role of decoding in working memory error reduction, and encourages future study of the decoding phase in more complex memory tasks.

Beyond memory error reduction mechanisms, how RNNs adapt to the entire environmental prior is an interesting problem. Adaptation is not simply about reducing the memory error for all colors. As observed in Figure 1D, also in previous works^1,9,10^, some low-prior colors exhibit larger memory errors. Thus, there is a trade-off for the RNN: how much error should be reduced for high-prior colors and how much should be increased for low-prior colors. This trade-off is unavoidable due to the neural mechanism. Introducing attractors to reduce memory error for common colors unavoidably leads to biased (or truncated) errors for nearby colors^1,10^. In the angular occupancy (Figure 3) part, enlarging common color’s angular occupancy will lead to smaller occupancy for other colors. How do RNNs balance memory errors among different colors remains unclear. Possible explanations can come from the Bayesian inference framework^21,53^. For example, Sohn et al.^36^ assumes that the agent adapts to the environmental prior by making Bayesian inference in each trial. Environmental prior is the prior function, noise distribution is the likelihood function. Based on the prior and likelihood, they proposed that the agent needs to output a memorized information value (in that paper was time duration) which equals to the mean of the posterior distribution. This theory can make concrete prediction on the neural dynamics. Alternative theory suggests the network can be a generative model for reproducing the environmental prior^17^. RNN trained in this paper provides a ready-to-use models for theory pre-testing.

## Materials and Methods

### RNN architecture

The RNN consists of 12 perception input neurons, 1 go input neuron, 12 response output neurons and 256 fully connected recurrent neurons (Fig. 1A). The dynamic equation of the RNN is^54^

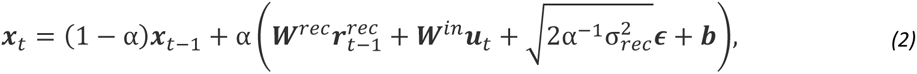

where ***x***_*t*_ is a 256-dimensional vector representing the activities/state of recurrent neurons (neural state) at time step *t*, and *α* = Δ*t* / τ, where Δ*t* is the time length for each computational step, *τ* is the neuron membrane time constant. In this study, we set Δ*t* = 20 ms and *α* = 1, which is the same choice as in previous work^25,54,55^. The change of neural state *x*_*t*+1_ subjects to 4 factors: (1) The inputs from the other recurrent neurons, 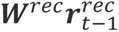, where 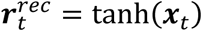 and ***W**^rec^* is the recurrent weights but the diagonal elements fixed to 0 (self-connections are not allowed)^54^ (2) The inputs from perception and go neurons, ***W***^*in*^***u***_*t*_, where ***u***_*t*_ consists of 12 components for the perception neurons and 1 component for the go neuron, and ***W***^*in*^ is the input weights. During the perception epoch, the perception neurons is entangled with an addictive gaussian noise with standard deviation *σ*_*x*_ = 0.2. (3) The intrinsic recurrent noise in the RNN, 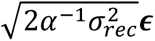, where ***ϵ*** is a 256-dimensional noisy vector drawn from the standard normal distribution at each time step, and *σ_*r*e*c*_* is the recurrent neural intrinsic noise strength. *σ_*r*e*c*_* = 0.2 was used in all main figures, except in Figure 5 where we inspected different noise strengths. (4) A constant bias current vector ***b***.

The activities of the 12 response neurons are read from the 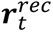,

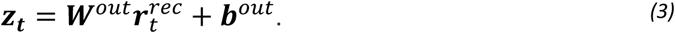

### A delayed-response trial and RNN training

A trial contained a fixation, perception, delay, go, and response epochs. During the fixation epoch, the RNN didn’t receive any inputs (i.e., ***u***_*t*_ = 0). In the perception epoch, a color *ϕ*_*s*_ was drawn from a prior distribution,

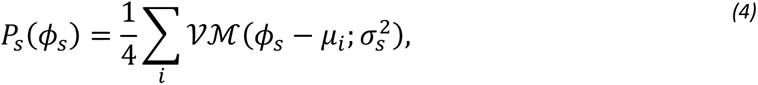

where **V*M*(⋅) is a von Mises function (circular version of the normal distribution) with mean values at *μ*_*i*_ ∈ {40^∘^, 130^∘^, 220^∘^, 310^∘^}

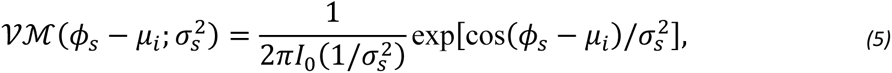

where σ_s_ is a free parameter to control the “width” of the prior distribution, and *I*_0_(·) is the modified Bessel function of order 0 for normalization purpose. Units of *ϕ*_*s*_, *μ* and *σ*_*s*_ were converted from degree to rad before feeding into the von Mises function. A color was drawn from the above prior distribution and then was sensed by the 12 perception neurons. Perception neurons had tuning curves that were also von Mises functions 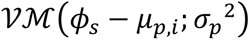 with 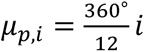 and 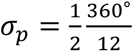 for *i* = 0, 1, …, 11 (Fig. 1A). Therefore, in the perception epoch, the perception components of the input vector ***u***_*t*_ are 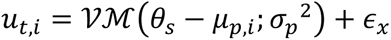 where *∈*_*x*_ is a gaussian noise with standard deviation *σ*_*x*_ = 0.2, and the go component is 0 (*u*_*t*,12_ = 0).

Next, during the delay epoch, all inputs were 0. The delay length was drawn from an uniform distribution *U*(0 ms, 1000 ms) in each trial. Then, in the go epoch, we set *u*_*t*,12_ = 1 to indicate a go cue and the rest of inputs and the outputs were 0. Finally, in the response epoch, the 12 response neurons were required to reproduce the neural activity of same as the perception neurons during their perception epoch, i.e. 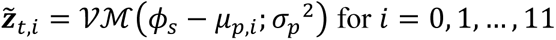.

The loss function is

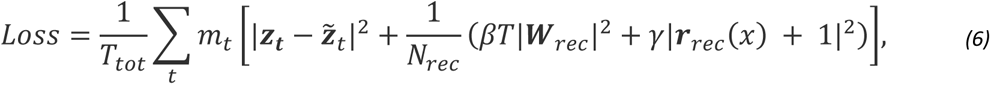

where *m*_*t*_ is the mask that is 0 in the fixation epoch and 1 in the rest epochs, *T*_*tot*_ is the trial time length, *N_rec_* is the number of recurrent neurons, *β* and *γ* in the regularization terms constrain the recurrent weights and neural firing rates. ***z̃**_t_* is the target response output. It was zero across perception, delay and go epochs, and was the target response 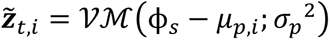 during the response epoch. This loss function was minimized by gradient descent using the Adams optimizer^56^.

Directly training the RNN to optimize the above loss function is difficult, we employed the progressive training protocol^27^: First, we trained the RNN with no noise, no regularization terms, uniform prior distribution, and delay epoch length fixed to 0. Secondly, we retrained the RNN with delay epoch length drawn from *U*(0 ms, 1000 ms). Thirdly, the RNN is retrained considering the noise and regularizations. The resulting model is called the pretrained RNN. Finally, we retrained the pretrained RNN with the desired prior distribution. If the desired prior distribution is still a uniform prior, then the trained RNN was called as Uniform RNN (approximately equivalent to the prior σ_*s*_ = 90^∘^), otherwise was called as Biased RNN.

### Mapping response neural activity into color

The actual response neural activity can be mapped back to a color by a population method. The mapped color is also called the RNN’s output color. More precisely, the response neural activity ***z***_***t***_ was firstly taking averaged within middle time window of the response epoch (60 ms to 140 ms, note the total response epoch is 200 ms). Next, the averaged response neural activity ***z̅*** was converted into the output color by a populational vector method^54,57^

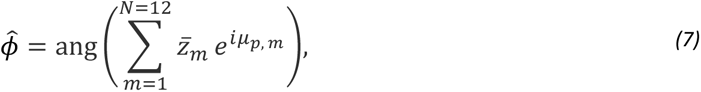

where ang(⋅) means taking the angle of a complex number, *i* here is a unit imaginary number, and *z̅*_*m*_ is the *m*th components of vector ***z̅***, i.e. the *m*th response neuron. 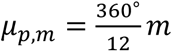 which was set to be align with the perception tuning mean.

### Removing outlier RNNs

Due to the stochastic nature of the training process, some RNNs exhibited significantly different metric values (e.g., memory error, entropy etc.) compared to others, this may be due to a failure of training. To mitigate the effect of these outliers, this paper employs a criterion for exclusion: any RNN whose measured metric exceeds 1.5 times the interquartile range (IQR) above the third quartile or below the first quartile is removed from the analysis. This exclusion was specified by the phrase ‘outlier RNNs were removed’ in figure legends.

### Identifying fix points

The dynamic equation of the RNN, Eq. (1), can be reformulated in the form of

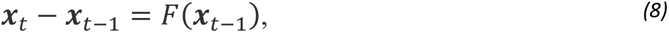

which defines the velocity field ***𝒗***(***x***) = *F*(***x***) / Δ*t*. In this study, we set Δ*t* = 20 ms. The global minimum points of speed *𝑣*(***x***) = |***𝒗***(***x***)| = 0 are the fixed points. Specifically, we ran the trained RNN with 500 trials with stimuli equally distributed from 0^∘^ to 360^∘^. The neural states at the beginning of delay epochs were collected. According to the visualization in Figure 3, these neural states were likely to near the ring on the delay. Hence these neural states were used as the initialization of searching point. Starting from these initial neural states, we searched for the global minima of speed using the gradient descent, where the independent variable is the position ***x***. Due to the imperfection of the gradient descent algorithm, we may end up with local minima. Nevertheless, these local minimum points are similar to the fixed points^58^, hence they are also called fixed points in this paper. For each fixed point, we computed the eigenvalues and eigenvectors of its Jacobian matrix 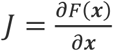. A fixed point is called an attractor if its largest eigenvalue is negative, while it is called saddle point if some eigenvalues are positive. The eigenvectors of the positive eigenvalues are projected and shown on the PC1-PC2 plane (Fig. 3). Only fixed points near the delay ring were shown.

### Decoding an end-of-delay neural state

We built an RNN decoder to decode a delay neural state 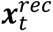 to its corresponding color. This RNN decoder faithfully represents the decoding phase of the RNN. Specifically, an RNN decoder is the same as the original trained RNN, but its recurrent neural state is initialized as the to-be-decoded neural state. Then, the RNN decoder runs directly through the go and response epoch. Response neural activities in the response epoch were temporally averaged. Averaged response was then decoded to an output color by the population vector method (Equation 7). This RNN decoder was used, for example, to decode attractors color (Figure 3D).

We also used the RNN decoder to decode the PC1-PC2 plane of the delay. A RNN was firstly ran through 1000 trials with uniformly distributed input color. Neural states during the delay were collected, used for fitting the PC1-PC2 plane, also called the delay plane. Next, a grid of mesh points (50 times 50) was sampled in the delay plane, centered at the delay plane center. Each of these mesh points were then PCA inversed back the original high-dimensional space, finally decoded by the RNN decoder (Figure 3G).

We also used the RNN decoder to find the relation between the angle of the delay plane and the color. Similar as above, a RNN was firstly ran through 1000 trials with uniformly distributed input color. Neural states at the end of delay (delay length is fixed as 800 ms) were collected, for fitting the PC1-PC2 plane (delay plane). Next, within the delay plane, a radius was computed, which was defined as the averaging distances of all collected 1000 neural states to the delay plane center. Using this radius, dense points (1000 points) of a ring were resampled. Finally, each of the resample points were decoded by the RNN decoder. This provides a mapping from the angle of the delay plane to the color (Figure 3H).

### Cross-decoding experiment

Cross-decoding involves preparing delay neural states using one RNN model (model 1) and decoding the prepared delay neural state using another RNN model (model 2). Specifically, model 1 was randomly chosen from either the Biased RNN or Uniform RNN categories. This model underwent 500 trials with a fixed input of a common color. Neural states at the end of delay were collected. Next, these end-of-delay neural states were used to set the recurrent neural states of model 2 which was randomly selected from the second category (which may be the same as model 1’s category). Model 2 processed these states over its decoding phase, outputted colors. The squared-root of the mean squared error across 500 trials was defined as the memory error. This procedure was repeated to gather memory error data for 50 pairs from each category combination, namely Bias & Bias, Bias & Uniform, Uniform & Bias, and Uniform & Uniform. The first category represents model 1 while the second represents model 2.

A key challenge in this process is the “neuron-matching problem” which involves determining how to match neurons in model 2 with those in model 1. Our approach to neuron matching is based on comparing the neural preferred color rank. The preferred color for a neuron in an RNN model is determined as the following. Dense neural states along a ring on the PC1-PC2 delay plane were sampled; then were decoded into output colors. This provides a delay-neural-states-to-color mapping. Since delay neural state is nothing but a vector of concatenated individual neural activations, delay-neural-states-to-color mapping was also serves as a neural-activation-to-color mapping. The preferred color for each neuron was defined as the color that maximized its neural activation.

We computed the preferred colors of all recurrent neurons in Model 1 and ranked them from 0 degrees (smallest) to 360 degrees (largest). We repeated this process for Model 2. We achieved neuron pairing by matching these ranks. Therefore, the prepared delay neuron state in Model 1, which includes 256 neurons, can correspond one-to-one with the recurrent neural state in Model 2, which also comprises 256 neurons. Subsequently, model 2 underwent the decoding phase to compute the memory error.

### A Taylor expansion approximation to the memory error

The squared memory error of a fixed common color *ϕ*_*c*_ is

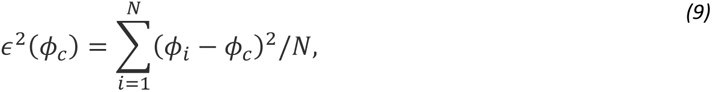

where *N* is the number of trials, *ϕ*_*i*_ is the RNN’s reported color in trial *i*. Approximately the RNN encodes a color by the angle (θ) of the delay plane, so output color is a function of the neural state’s angle *ϕ*(*θ*). Denoting the angle encoding common color as *θ*_*c*_. Assuming during delay the actual neural state’s angle doesn’t deviate from *θ*_*c*_ too much, then we can Taylor expand 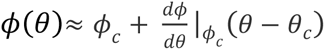. Substituting this back to Equation (9) yields

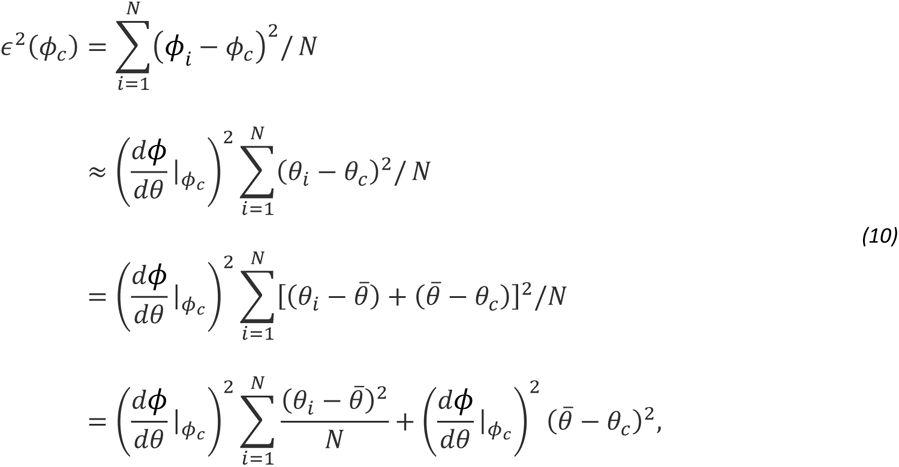

were *dϕ*/*dθ*|_*ϕc*_ is the reciprocal of the angular occupancy at common color, 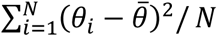 is the dynamic dispersion, and the last term describes a correction from the actually neural state mean angle to the common color angle.

### Numerical procedure for testing the Taylor expansion approximation

To test this approximation formula (Equation 10), we computed the dynamic dispersion, angular occupancy and mean bias correction, respectively. To compute the dynamic dispersion, we ran each of the RNN 5000 trials with fixed input color as common color (delay length 800 ms). Neural states at the end of the delay were collected, and their corresponding angles *θ*_*i*_ were computed. Outlier angles (1.5 IQR below the first quantile or above the third quantile) were removed. Dynamic dispersion was measured as the variance of *θ*_*i*_. The mean of *θ*_*i*_ was also denoted as *θ̅*. To compute angular occupancy, similar as Figure 3H (see Methods), dense points along a ring were sampled in the delay plane. Each of these points was decoded by the RNN decoder, hence we obtained the angle-color mapping. Numerical differentiation was then performed to compute the angular occupancy. In addition, the angle on the ring whose color was closest to the common color was denoted as *θ*_*c*_. Finally, using the *θ̅* and *θ*_*c*_ computed previously, mean bias correction can be computed as written in the Equation (10).

The resulting memory error computed from all these terms (equation 10) was called theoretical prediction. To explore the importance of decoding strategy (i.e. angular occupancy), as a comparison, we also computed theoretical prediction but setting angular occupancy to 1.

For the experimental RNN’s memory error, for each condition (e.g. environmental prior σ_*s*_ or noise level σ*_rec_*), 50 RNNs were used. Each RNN was ran on 5000 trials. After removing the outliers’ trial errors, each RNN’s experimental memory error is the trial error’s standard deviation (squared root of Equation 9).

### Decoding neural states on the response plane

A neural state on the delay plane can decoded into an output color by using the RNN decoder which is continuing running the RNN through go and response epochs. Similarly, a neural state in the response plane can also be decoded into an output color by continuing running the RNN. Since the response is already the last epoch, hence the “continuously running” only means reading the (recurrent) neural state out to the response neuron activity, by using RNN’s readout weight ***W^out^*** and bias current ***b^out^*** (Equation 3). The response neuron activity was then translated into an output color using populational vector method (equation 7). Note that no dynamics is involved since no time component here. This decoding procedure is also called readout decoder. Readout decoder’s properties is fully determined by the readout weight and bias currents.

To inspect the properties of Readout process, we used the Readout decoder to decode mesh points on the response PC1-PC2 plane. Specifically, each RNN was run through 1000 trials, neural states during the response epochs were collected for fitting the response plane (PC1-PC2 plane). After fitting, we mesh grid points on the response plane. They were further decoded by the Readout decoder.

Similarly, we can also inspect the relation between angle and the color in the response plane. The procedure is analogous to what was described in the “Decoding an end-of-delay neural state” session, except that the delay plane was replaced by the response plane, and readout decoders were used instead of RNN decoders.

## Acknowledgments

We thank Daoyun Ji from the Baylor College of Medicine for constructive comments and suggestions to this work.

## Funding

Hong Kong Baptist University Strategic Development Fund

the Hong Kong Research Grant Council (GRF12200620)

the Hong Kong Baptist University Research Committee Interdisciplinary Research

Clusters Matching Scheme 2018/19 (RC-IRCMs/18 19/SCI01)

the National Science Foundation of China (Grant 11975194)

## Author contributions

Conceptualization: ZY, LT, CZ

Methodology: ZY, HL, LT, CZ

Investigation: ZY, HL

Supervision: LT, CZ

Writing—original draft: ZY, HL, LT, CZ

Writing—review & editing: ZY, HL, LT, CZ

## Competing interests

Authors declare that they have no competing interests

## Declaration of generative AI and AI-assisted technologies in the writing process

During the preparation of this work the author(s) used ChatGPT-4 in order to polish the language and improve the readability of the manuscript. After using this tool/service, the authors reviewed and edited the content as needed and take full responsibility for the content of the published article.

## Data and materials availability

The codes for training and analysis are available at https://github.com/AgeYY/working_memory.git

## Supplementary Materials

**SI Figure 1.**
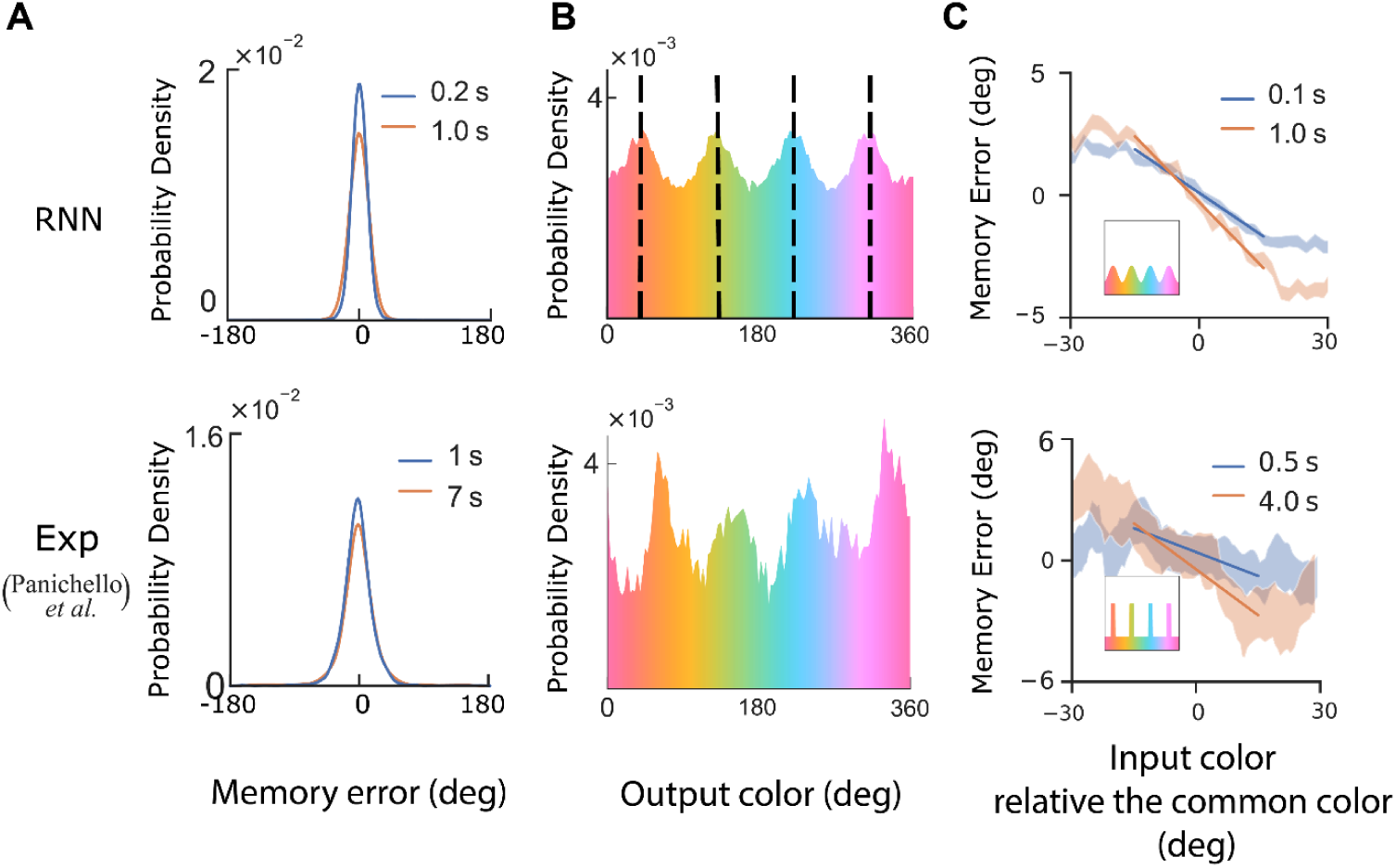
Trained Biased RNN (σ_*s*_ = 25°) exhibited behavior features similar to the experiments. (**A**) The distribution of memory error. Each RNN was tested on 1000 trials with uniformly distributed input color. Memory error from 50 Biased RNNs were concatenated. Bottom: experimental data. (**B**) Upper: the distribution of output color when the input colors were sampled uniformly for each trial. 50 Biased RNNs results were concatenated, 1000 trials each. Black dashed lines indicate common colors in the training prior distribution. Bottom: Human’s output color distribution. Note human had not yet been trained on biased prior color distribution. The biasness observed here may come from the biased color prior in the natural environment. (**C**) Upper: memory error for each input color. 50 Biased RNN was used, each input color was run through 1000 trials for each of the Biased RNN. The error band is the standard error of the mean across trials and RNNs. The delay lengths were fixed to either 0.1s or 1s. Inset shows the training environmental prior. Bottom: human participants were trained on a biased prior distribution (inset). Their performance during the last third of the training trials is shown. Color represents results when delay was fixed to either 0.5s or 4.0s. Experimental figures were plotted using the data published by Panichello et al^1^.

**SI Figure 2.**
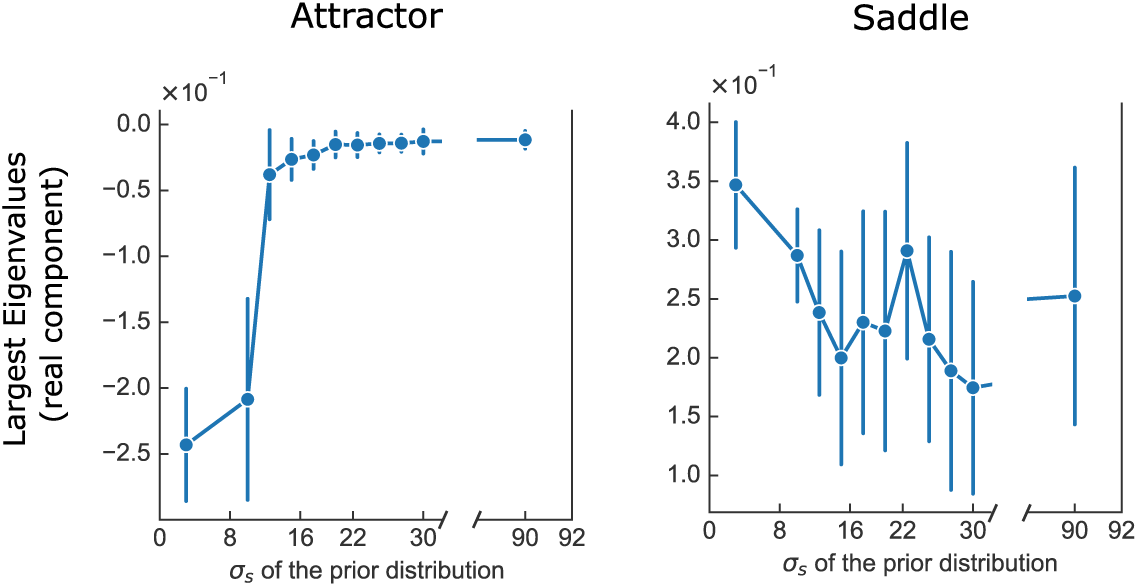
Eigenvalues of the fixed points. (Left) We trained 50 RNNs for each prior distribution. For each RNN, we searched for the attractors (see Methods). The largest eigenvalue (real component) for each attractor is denoted as *𝑣*_*i,j*_, where *i* indicates the *i*th RNN and *j* indicates the *j*th attractor. The averaged largest eigenvalues across different attractors within the same RNN is denoted as 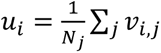. Dots in the figure represent the mean of largest eigenvalues across different RNNs, 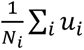. Error bars indicate the standard deviation of *u*_*i*_ across different RNNs. (Right) Same as left but considering the largest real component of saddles. Both attractors and saddles become stronger (larger absolute values for the real parts) for narrower prior distributions.

**SI Figure 3.**
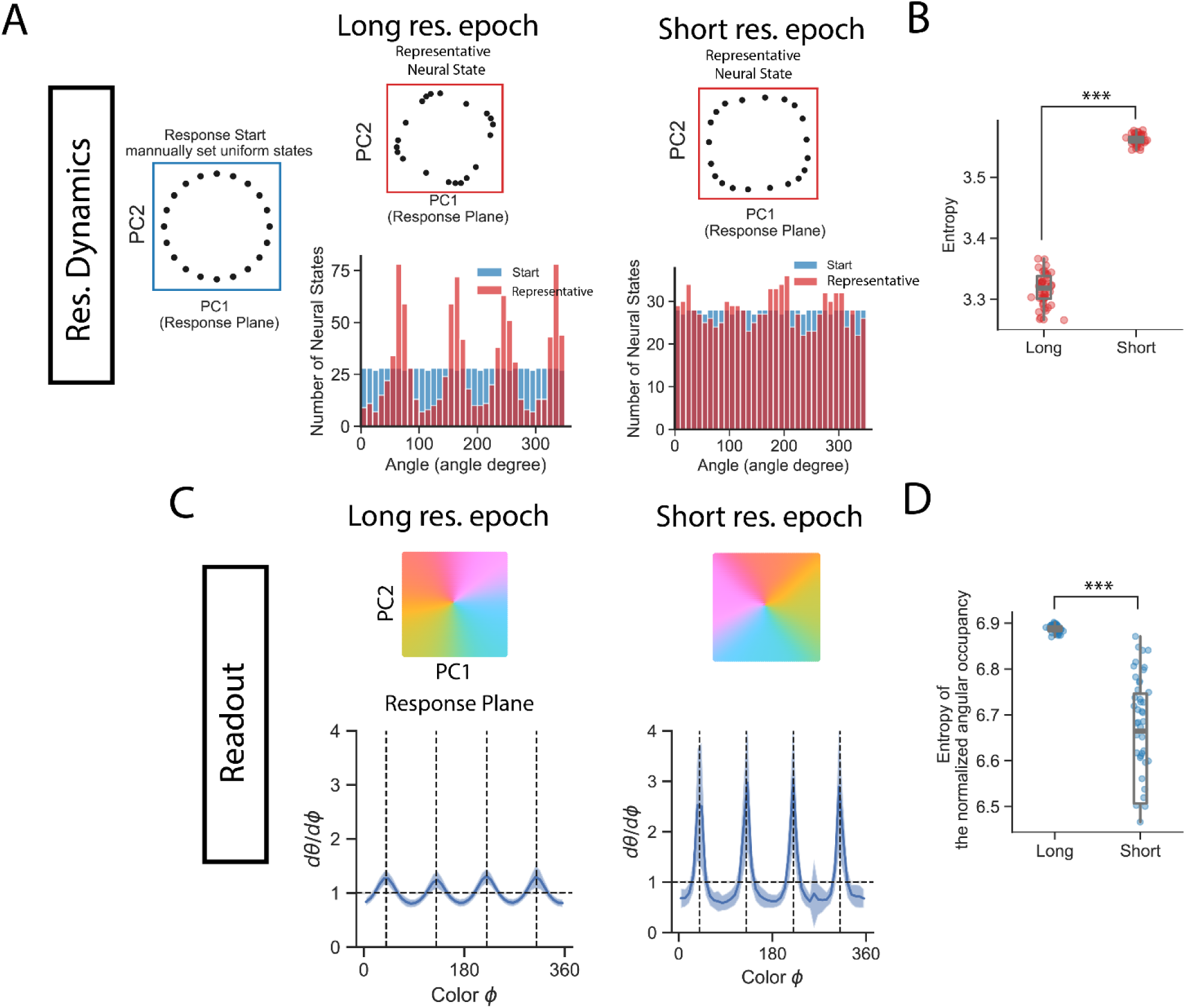
RNNs trained on a shorter response epoch have weaker response dynamics bias and stronger readout bias. Training prior *σ*_*s*_ = 3^∘^. Computing details are identical to Figure 6 except that the RNN models were replaced by which trained with long/short response (res.) epoch. (**A**) Long/short response epoch: one example RNN trained with long (200 ms)/short (40 ms) response epoch duration. Compared to the RNN trained on long response epoch, RNN trained on short response has smaller change of neural states during the response epoch. (**B**) Increasing entropy of the representative neural state distribution also suggests that RNNs trained on short response epoch have less neural states’ angular changes during the response. Each dot is one RNN’s representative neural state entropy (red histogram in panel A). ***: *p* < 10^−3^ Wilcoxon signed-rank test. (**C**) RNNs trained on short response epoch have more biased readout. Line shows the mean of 50 RNNs, and error band is the standard deviation. Dash lines: four common colors. Outlier RNNs were removed. (**D**) Entropy of angular occupancy, dot is an entropy of one RNN’s angular occupancy (normalized). Outliers were not shown. ***: *p* < 10^−3^ Wilcoxon signed-rank test.

## Notes

### Competing Interest Statement

The authors have declared no competing interest.

### Summary of Updates

Merging figure one and two; polishing language

